# Oxygen Transport in Nanoporous SiN Membrane Compared to PDMS and Polypropylene for Microfluidic ECMO

**DOI:** 10.1101/2025.01.04.631337

**Authors:** Nayeem Imtiaz, William A. Stoddard, Steven W. Day

## Abstract

Extracorporeal Membrane Oxygenation (ECMO) serves as a crucial intervention for patients with severe pulmonary dysfunction by facilitating oxygenation and carbon dioxide removal. While traditional ECMO systems are effective, their large priming volumes and significant blood-contacting surface areas can lead to complications, particularly in neonates and pediatric patients. Microfluidic ECMO systems offer a promising alternative by miniaturizing the ECMO technology, reducing blood volume requirements, and minimizing device surface area to improve safety and efficiency. This study investigates the oxygen transport performance of three membrane types— polydimethylsiloxane (PDMS), polypropylene, and a novel nanoporous silicon nitride (SiN) membrane—in a microfluidic ECMO platform. While nanoporous membranes rely on pore-mediated diffusion and PDMS on polymer lattice diffusion, results show no significant differences in device oxygenation efficiency (p > 0.05). Blood-side factors, including the diffusion rate of oxygen through the red blood cell (RBC) membrane, RBC residence time, and hemoglobin binding kinetics, were identified as primary bottlenecks. Even computational models of a hypothetical infinitely permeable membrane corroborate the limited impact of membrane material. These findings suggest a shift in ECMO design priorities from membrane material to blood-side enhancements. This research provides a foundation for optimizing ECMO systems.

---

1. Introduction

The prevalence in chronic lung diseases, such as Chronic Obstructive Pulmonary Disease (COPD), along with sudden outbreaks of infectious diseases like swine flu and COVID-19, has highlighted the need for better treatments for respiratory insufficiency and respiratory failure (Berlin et al. 2020; Roberts et al. 2024). Mechanical ventilation is the standard treatment, but it involves invasive procedures that carry serious risks, such as barotrauma, ventilator-associated pneumonia, and other infections (Fan et al. 2017). In response, Extracorporeal Membrane Oxygenation (ECMO) has become increasingly significant. ECMO is a technique that circulates blood through an external circuit to allow gas exchange in an artificial lung, which can help reduce or, in some cases, avoid the need for a ventilator (Auld et al. 2020; Li et al. 2020).

Although ECMO has become safer and more effective over the past 20 years, there are still important areas that need improvement. Patients on ECMO are at risk for several complications, including infection, thrombosis, embolism, and hemorrhage (Aubron et al. 2013).

Microfluidic-based ECMO devices have the potential to transform ECMO treatment by significantly reducing blood volume, blood-contacting surface area, and overall device size. This reduction in blood volume benefits patients of all ages and is especially crucial for smaller patients, such as neonates. Additionally, minimizing the membrane and device surface area, lowers the risk of complications and reduces blood damage (Dabaghi et al. 2018a; Rode et al. 2020). Current oxygenators usually have a tube-in-tube design (Odish et al. 2023; Shin et al. 2024), whereas many microfluidic devices, such as lab-on-a-chip systems, are arranged in stacked planar layers (Abudupataer et al. 2020; Hou et al. 2018). This stacked configuration is compatible with a new generation of ultra-high permeability membranes that are manufactured on a wafer and must remain planar (Dabaghi et al. 2018a; Rode et al. 2020) Various groups have made significant progress in the development of microfluidic oxygenators over the past decade, using advances in computational designs, microfabrication techniques, and biomaterials technologies to create prototype devices that have been tested *in vitro* and in numerous proof of concept studies (Abada et al. 2018; Burgess et al. 2009; Dabaghi et al. 2018b; Imtiaz et al. 2023; Imtiaz et al. 2022).

Regardless of great advancements in microfluidic ECMO research, challenges still remain in reducing priming volume and membrane surface area, particularly for neonatal and pediatric applications. Developing advanced membrane and device technologies to create miniaturized ECMO systems with lower priming volumes is therefore essential. The majority of devices in the literature utilize PDMS as the membrane material (Dabaghi et al. 2019; Ma et al. 2022; Potkay 2014; Santos et al. 2021), leaving the potential benefits of more advanced, novel membrane materials largely unexplored. It has been proposed that using a high gas-permeant membranes enable greater oxygen transfer across the membrane surface, which could result in more efficient oxygen delivery over time (Gimbel et al. 2021), eventually resulting in reduced device size.

To miniaturize the microfluidic ECMO it is important to select a membrane with high gas permeance. High gas-permeant membranes enable greater oxygen transfer across the membrane surface, which could result in more efficient oxygen delivery over time (Gimbel et al. 2021).

PDMS membranes used in current microfluidic ECMO devices offer limited gas-permeance. Conventional ECMOs, on the other hand, typically employ polypropylene membranes, which, while more permeable than PDMS, are still not a significant improvement.

Nanomembranes hold significant promise for transforming a wide range of fields, including separation processes, energy production, medical applications (Blauvelt et al. 2023; Conlisk et al. 2009; Desai et al. 1999; Gimi et al. 2009; Han et al. 2008a; Haque et al. 2013; Jeon et al. 2012; Kim et al. 2008; Qi et al. 2014; Siwy et al. 2005; Snyder et al. 2011). These membranes have since been utilized in numerous applications, such as cell culture, electro-osmotic pumping, hemodialysis, lab-on-a-chip devices, and investigations into portable hemodialysis systems (Han et al. 2008b; Hill et al. 2020; Stroeve and Ileri 2011). The development of nanoporous SiN membranes has inspired their exploration of microfluidic ECMO applications. The permeability of nanoporous SiN is not only three orders of magnitude higher than that of polydimethylsiloxane (PDMS), but also, due to its ultrathin nature (<500 nm), the gas-permeance of the nanoporous SiN membrane is five orders of magnitude higher than PDMS (Formica et al. 2008; Miller et al. 2020). The membrane thickness, gas permeance, permeability, and membrane resistance values for the membranes investigated in this study are summarized in Table 1.

**Table 1.** Membrane thickness, gas permeance, permeability, and membrane resistance information for the nanoporous SiN, PDMS, and Polypropylene membrane.

| Membrane Material | Permeability ($P$) | | Membrane Thickness ($L$) | Gas Permeance ($P_m$) | | Membrane Resistance ($R_m$) | |
| --- | --- | --- | --- | --- | --- | --- | --- |
|  | (cc·cm·min<sup>-1</sup>·bar<sup>-1</sup>·cm<sup>-2</sup>) | m<sup>2</sup>/(s·Pa) | (μm) | (cc·min<sup>-1</sup>·bar<sup>-1</sup>·cm<sup>-2</sup>) | m/(s·Pa) | (min·bar·cm<sup>-1</sup>) | (s/m)·Pa |
| Nanoporous SiN | 0.54 | $9.0 \times 10^{-12}$ | 0.4 | 13,500 | $2.3 \times 10^{-5}$ | $7.4 \times 10^{-5}$ | 44,400 |
| PDMS | $2.7 \times 10^{-3}$ | $4.5 \times 10^{-14}$ | 20 | 1.4 | $2.3 \times 10^{-9}$ | 0.72 | $4.3 \times 10^8$ |
| Polypropylene | 76.5 | $1.3 \times 10^{-9}$ | 170 | 4500 | $7.6 \times 10^{-6}$ | $2.2 \times 10^{-4}$ | 132,000 |

Permeability *(P)* is an intrinsic membrane property, gas permeance *(P_m_)* is Permeability divided by membrane thickness *(L), P_m_ = P/L,* membrane resistance *(R_m_)* is proportional to membrane thickness and inversely proportional to permeability, *R_m_ = L/P*.

Despite the potential of nanoporous SiN as an ultra-high permeability membrane, a comparative analysis of different membrane materials within a standardized device platform remains largely unexplored. Nanoporous SiN membranes could offer a promising alternative for micro ECMO applications due to their unique properties (Miller et al. 2020). In this study, we investigate the oxygen transport performance of three distinct membrane materials—nanoporous SiN, PDMS, and polypropylene—within a prototype modular microfluidic ECMO device platform (Figure 1), using both empirical methods and computational fluid dynamics (CFD) simulations. This comparative analysis is designed to evaluate the efficacy of nanoporous SiN as a novel membrane material. Additionally, we include a hypothetical model of an infinitely permeable membrane to represent an ideal scenario for oxygen transport. This work aims to provide the medical device community with deeper insights into the impact of membrane material selection in the design of microfluidic ECMO systems and blood-gas exchangers in general.

**Fig. 1.**
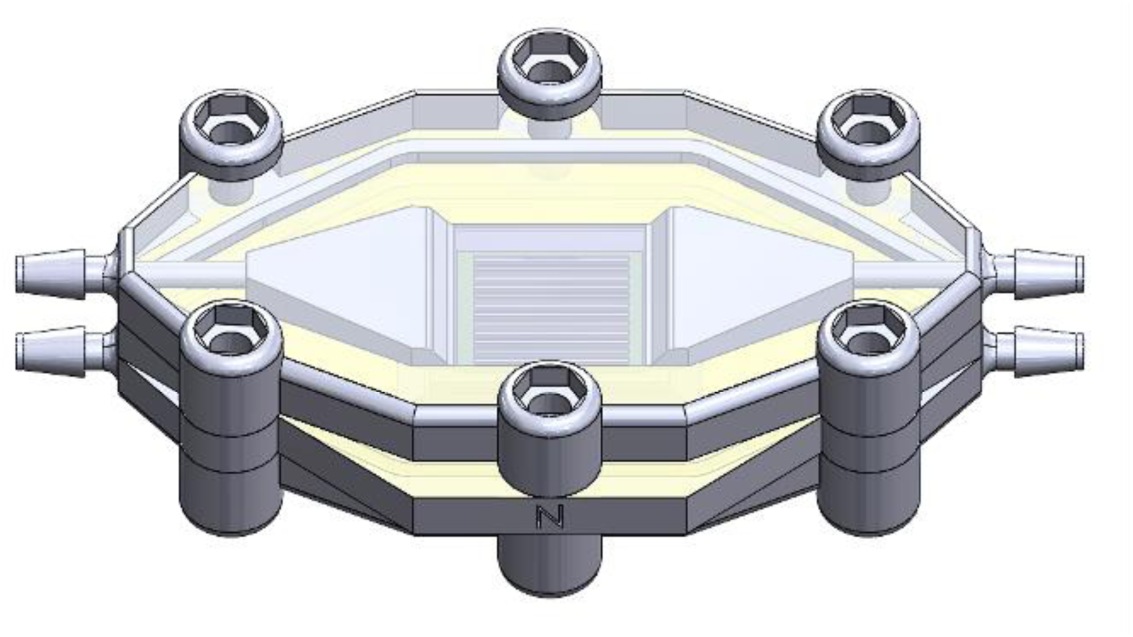
Modular microfluidic ECMO test device prototype

## 2. Material and Methods

### 2.1. Membranes

The nanoporous SiN membrane, with a thickness of 400 nm, 20% porosity, and 500 nm pore size, was obtained from SiMPore (NY, USA) (Hill et al. 2020). 20 µm thick PDMS membranes used in this study was also obtained from SiMPore, USA. The polypropylene membrane (Sterlitech (WA, USA)), had a thickness of 170 µm and nominal pore sizes of 200 nm.

### 2.2. Device Design and Fabrication

The microfluidic ECMO device prototypes were designed using SolidWorks 2020, and fabricated through 3D printing using a Form 3+ SLA printer (Formlabs) and biocompatible resin (BioMed Clear Resin). The device consists of two outer housings, a middle layer, and a membrane (Fig. 2a). A silicon frame was used to support the PDMS and polypropylene membranes, and frames were attached to the device’s middle layer with PDMS adhesive (Fig. 2a). The nanoporous SiN was sourced with the same silicon support frame as used for the PDMS and polypropylene membranes. After membrane attachment, the middle layer is sandwiched between two outer housings (Fig. 2a). The final assembly is the full microfluidic ECMO device prototype. The modular nature of the ECMO prototype allows for rapid exchange of the membrane layer. Fig. 2b shows the blood flow path through the blood side of the device. Table 2 shows the device dimensions in detail.

**Fig. 2.**
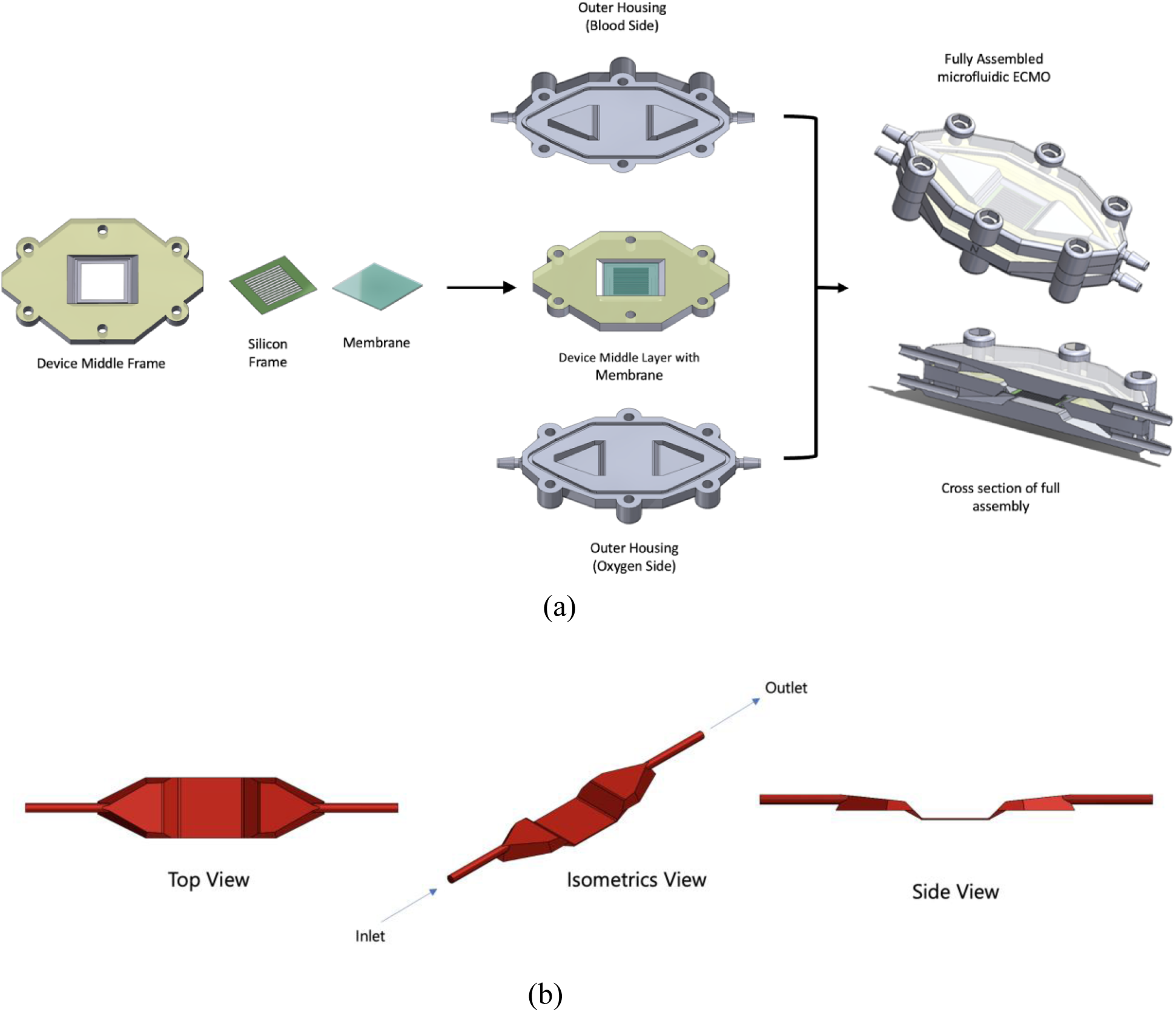
a) Full device assembly, with two outer housings and middle layer. b) Blood flow path in the blood side of the device

**Table 2.** Geometrical features of the prototype ECMO device.

| Property | Dimension |
| --- | --- |
| Inlet port inner diameter (mm) | 2.6 |
| Outlet port inner diameter (mm) | 2.6 |
| Device length (inlet to outlet) (cm) | 10.6 |
| Channel height (mm) | 0.4 |
| Active membrane area (cm<sup>2</sup>) | 1.77 |
| Device volume (cm<sup>3</sup>) | 1.73 |

### 2.3. Experimental Setup

The experimental setup is illustrated in Fig. 3. Two SpO₂ sensors (CritLine, USA) were placed at the inlet and outlet of the blood side of the device, labeled as S2 and S3, respectively, to monitor oxygenation levels. The pressure drop between the inlet and outlet of the blood side was monitored. O₂-saturated water was flowed at 8 ml/min on the gas side of the device. A peristaltic pump was used to recirculate blood through the loop, while a custom-designed blood reservoir served as both a reservoir and a dampener. To prevent atmospheric gas interference with the blood, the reservoir was equipped with a “floating boat” lid.

**Fig. 3.**
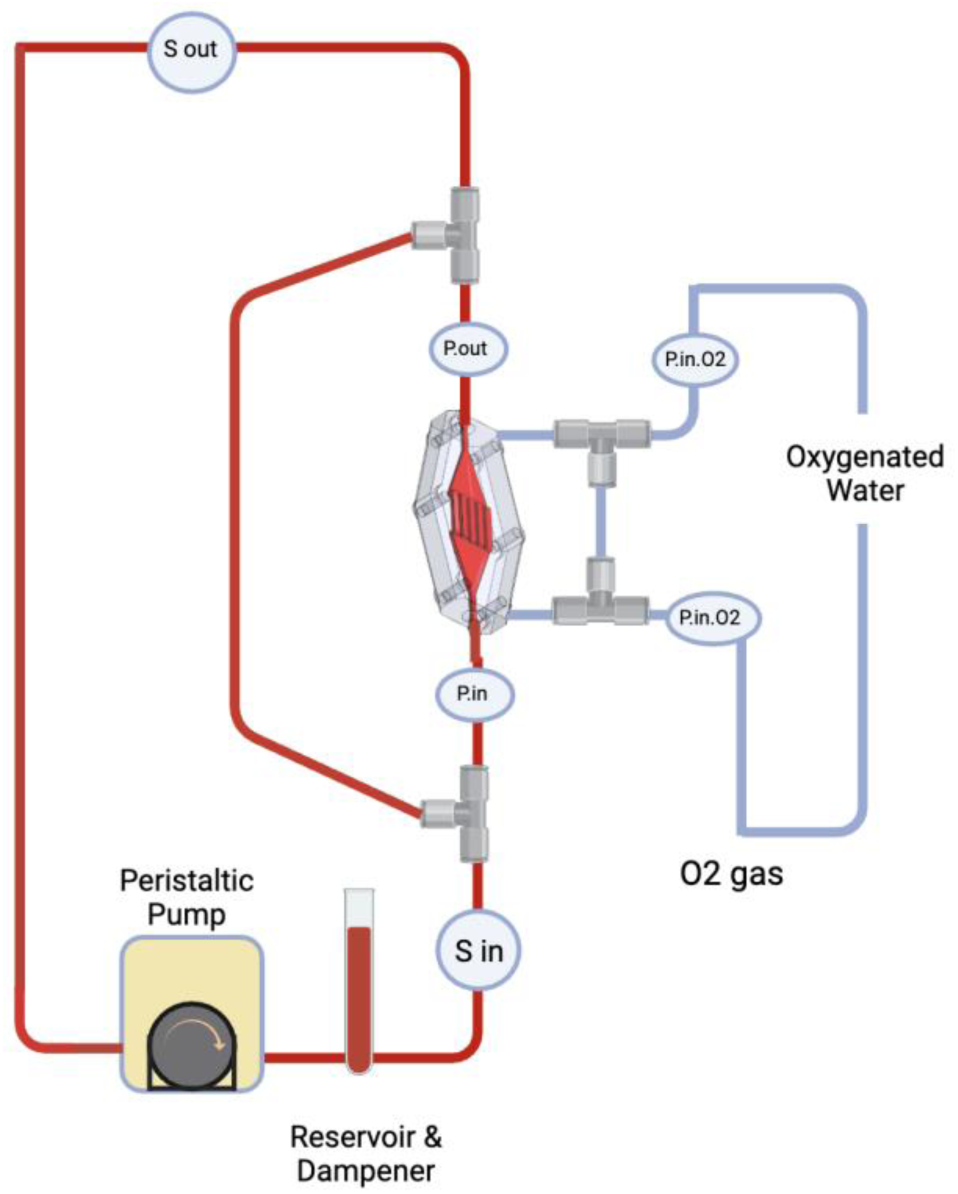
Oxygen transport in the blood through oxygenated water experimental setup

*Rationale for Using Oxygenated Water Instead of Pure Oxygen Gas:* The nanoporous SiN membrane is highly fragile, and exposure to pure oxygen gas on the gas side led to membrane failure, preventing the completion of experiments. To address this issue, oxygenated water was used as an alternative to pure oxygen gas. This approach provided a stable oxygen source without compromising membrane integrity, allowing the SiN membranes to remain intact throughout the experimental process.

Heparinized bovine calf blood (Lampire, USA) was reduced to an initial SpO₂ of 70% by bubbling the blood with nitrogen. A volume of 30 mL of prepared blood was then loaded into the continuous flow loop. Oxygen gain was measured and reported as SpO₂% increase per pass, SpO₂% increase per minute, mL O₂ gain per pass, and mL O₂ gain per minute. For each flow rate (0.2, 0.5, 1, and 2 mL/min) and membrane type, 4 to 6 samples (n = 4–6) were taken to ensure statistical robustness. The flow rate of the oxygenated water was 8 ml/min.

*Blood Viability Assessment:* A pre-experiment screening was conducted to ensure blood viability. In this assessment, the blood was bubbled with oxygen. Only blood that demonstrated an SpO₂ increase of at least 2% per minute in a 100 mL sample was considered viable.

### 2.4. Computational Model Setup

Our setup consisted of a device that flows blood through a channel adjacent to a membrane, with a second channel on the other side of the membrane, carrying oxygenated water. The geometry was generated from the files used to create the 3D-printed test article. The geometry for the flow path was imported to ANSYS Workbench 2020 for meshing. A hexahedral mesh was generated with refinements around areas of geometrical complexity. The highest refinement was in the section adjacent to the membrane. It had a target cell size of 0.04 mm. An inflation region at the membrane wall was implemented to capture the diffusion boundary layer accurately. The inflation zone is 10 cells thick, with a ratio of 1.2 per layer, resulting in the smallest cell height being 6 nm. The resulting mesh is 7.26 million nodes. Fig. 4 shows the Flowchart of the computational model for membrane oxygen transport.

**Fig. 4.**
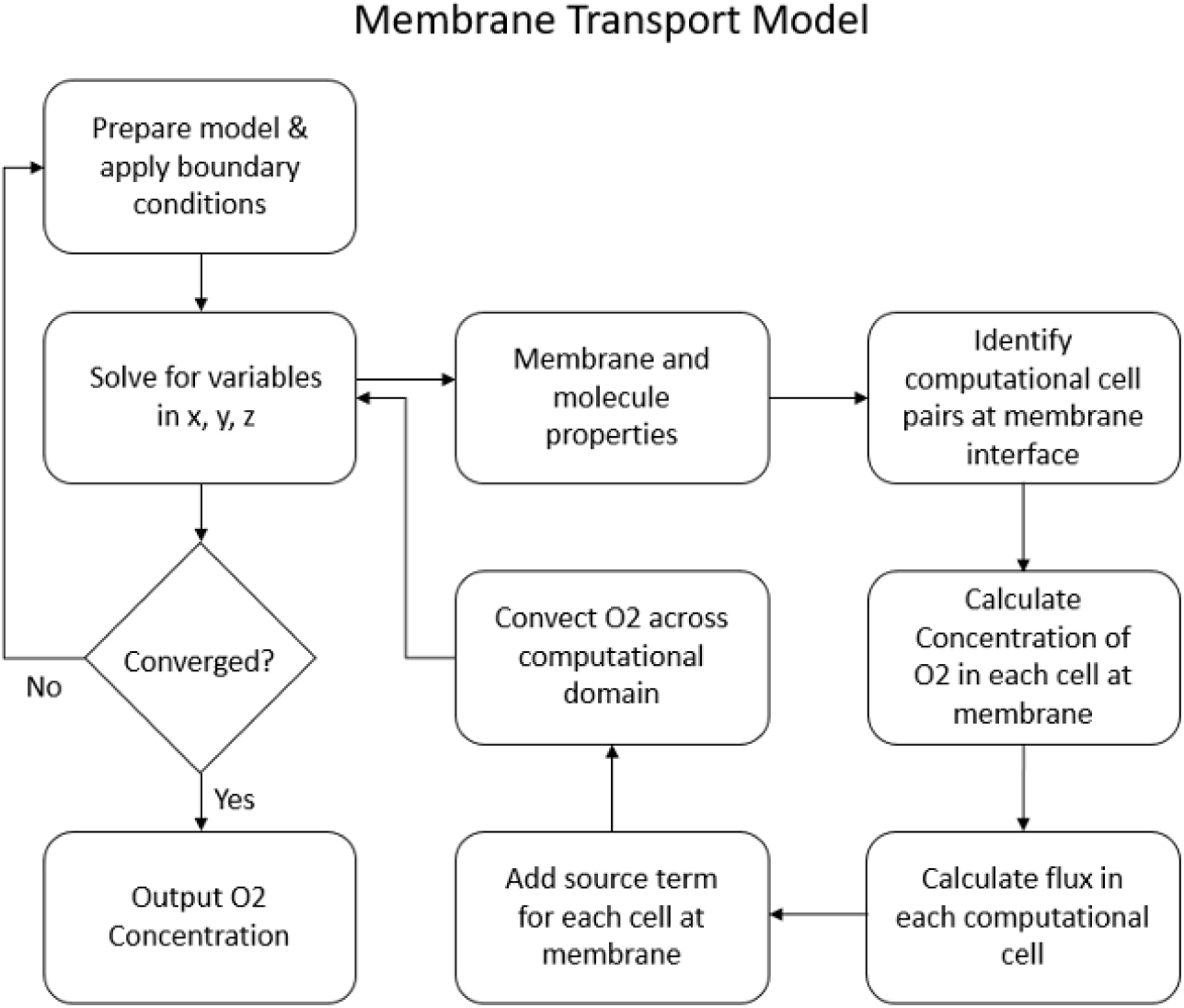
Flow chart showing the UDF process for oxygen transport across the membrane

The mesh was then imported into ANSYS Fluent 2020. Using computational fluid dynamics (CFD), ANSYS Fluent enables researchers to model complex blood flow behavior, including velocity profiles, shear stress, and pressure distributions within medical devices and blood vessels (Imtiaz et al. 2024; Nayeem Imtiaz and Tasfia Siam Siam 2023). A steady laminar flow solver was used to calculate the velocity field and transport. The oxygen transfer was simulated using a modified User Defined Function (UDF) code that originally simulated dialysis. The oxygen transport term was defined as a species and their transport was governed by the blood or dialysate flow fields. The oxygen-saturated water computational domain was mathematically coupled to the blood domain through the UDF architecture to implement source/sink and convective terms across the membrane, thus allowing species to move across the membrane.

The user-defined variable is in units of mg/L of O2. The concentration calculated for 70% saturation in human blood is entered as the blood side inlet. This matches the 70% oxygenation done in our bovine blood experiments for validation. The inlet for the oxygenated water side is initiated at 100% saturation for 1 atmosphere of O2. Henry’s law is used to calculate the concentration of oxygen in water (in mg/L) in the water side. Fick’s first law (Eq. 1) gives the diffusion across the membrane. The flux *J*i through an adjacent cell pair *i* at the membrane is calculated for each timestep, where *D*m represents the calculated membrane diffusivity and Δ*C* is the concentration gradient across the paired cells.

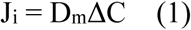

In order to account for the plasma oxygen concentration, a combination of the Hill Equation (Eq. 2) (Hill 1910) and Henry’s law for plasma are used to get a relationship between total blood oxygen content and the plasma concentration (Sové et al. 2016). An average hematocrit was chosen in the normal human range (Walker 1990). In Hill’s law (Eq.2), SO2 is the hemoglobin oxygen saturation, N is the Hill coefficient, PO2 is oxygen partial pressure and P50 is the partial pressure of oxygen at 50% saturation. The equations are not simple to solve, so a 3rd order polynomial trendline is used in the calculation as an approximation. The diffusivity of oxygen through a 20 μm PDMS membrane was compared to a SiN chip with cylindrical pores (Wright et al. 2020). The membrane resistance, which is the inverse of membrane permeability, was used in the model to dictate membrane properties (Table 1). Given an initial oxygen partial pressure difference (ΔP) of 120 mmHg, with the oxygenated water side at 100% O₂ saturation and the blood side at 70% O₂ saturation, the calculated membrane resistance was 2.78 s/m for SiN and 2688 s/m for PDMS. Additionally, a hypothetical infinitely permeable membrane (zero membrane resistance) was modeled to provide a “best-case scenario” and was compared to the PDMS and SiN membrane.

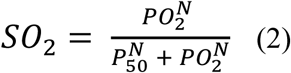

The mass flow of the blood is set for each case in the UDF, and a parabolic velocity profile is assigned to the inlet. The water flow is kept at a constant 8 ml/min to mimic the set-up of the experiment. Once the simulation has converged, a mass average of the outlet concentration of O2 is obtained. The initial concentration of O2 in plasma is subtracted from the outlet concentration. This results in the total concentration in mg/L that was transmitted into the blood. This is added to the 70% saturation total oxygen level of whole blood, including oxygen in the hemoglobin.

This is divided into the carrying capacity for the hemoglobin in blood to get the outlet saturation, SO2.

### 2.5. Data Analysis

The cross-membrane and cross-flow rate comparison for the empirical experiments was done using the one-way analysis of variance (ANOVA), where p < 0.05 indicates that two sets of data are distinct. The computational and empirical data for the membranes were compared by one-sample t-test, where a p < 0.05 indicates that the computational datum is distinct from the empirical data. Software tool Graphpad Prism 10 was used for data post-processing and statistics.

## 3. Results

### 3.1. Empirical Results

Fig. 5 presents the oxygen transport performance of the three membrane types across four flow rates. Fig. 6a illustrates the SpO₂% increase and ml O_2_ gain per pass for each membrane type as the full blood volume flows through the device. Results indicate that at any given flow rate, the O_2_ increase is not significantly different between membrane types (p > 0.05), where it is evident that per-pass oxygen transport decreases for all membranes as the flow rate increases. Fig. 5b display the SpO₂% increase and mL oxygen gain per minute. These data reveal that oxygen gain per minute does not differ significantly between membrane types at a given flow rate, and it increases for all membranes as the flow rate increases. The pressure drop between the inlet and outlet of the blood side remained within 0.03 to 0.2 PSI for all flow rates.

**Fig. 5.**
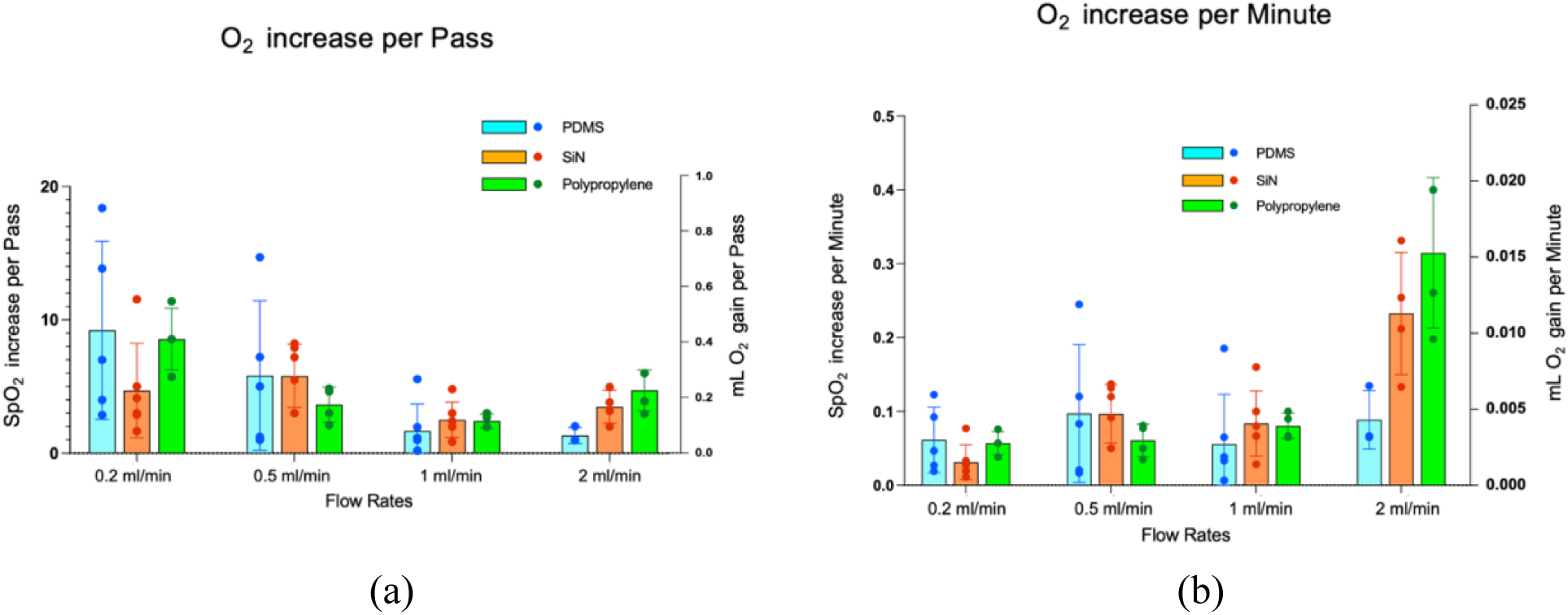
SpO2% rise and ml O_2_ gain of blood for PDMS, nanoporous SiN, and Polypropylene membrane at different flow rates, (a) per-pass, (b) per minute. In each panel, the left y-axis shows SpO_2_ % rise and the right y-axis shows ml O_2_ gain

**Fig. 6.**
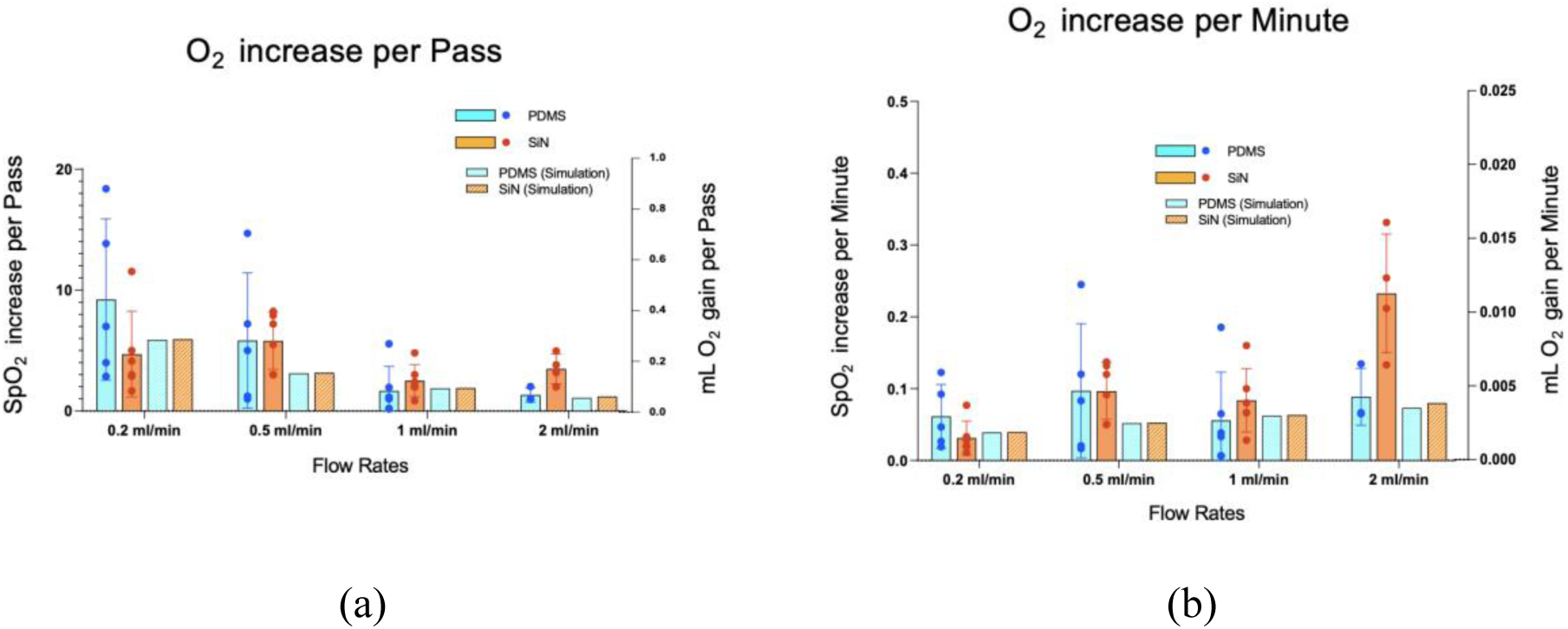
SpO2% rise and ml O_2_ gain of blood for PDMS and nanoporous SiN membrane at different flow rates (empirical and simulation), (a) per-pass, (b) per minute. In each panel, the left y-axis shows SpO_2_ % rise and the right y-axis shows ml O_2_ gain. (solid bar: empirical, patterned bar: simulation)

### 3.2. Empirical Vs. Simulation

Fig. 6 shows the oxygen transport comparison between the empirical and simulation data for PDMS and nanoporous SiN membrane. It is observed that for all flow rates the empirical agrees with the simulation results (p > 0.05), indicating that the simulation model was successful in predicting the oxygen transport. The oxygen transport behavior observed for the empirical described in Fig. 5 holds true for the simulations.

### 3.3. Hypothetical infinite permeability membrane (simulation)

Fig. 7 shows the computational oxygen transport values for the hypothetical infinite permeability membrane compared to the nanoporous SiN and PDMS simulations. The results indicate that the oxygen gain with the hypothetical perfect membrane is slightly higher than with the conventional membranes, but the increase is minimal.

**Fig. 7.**
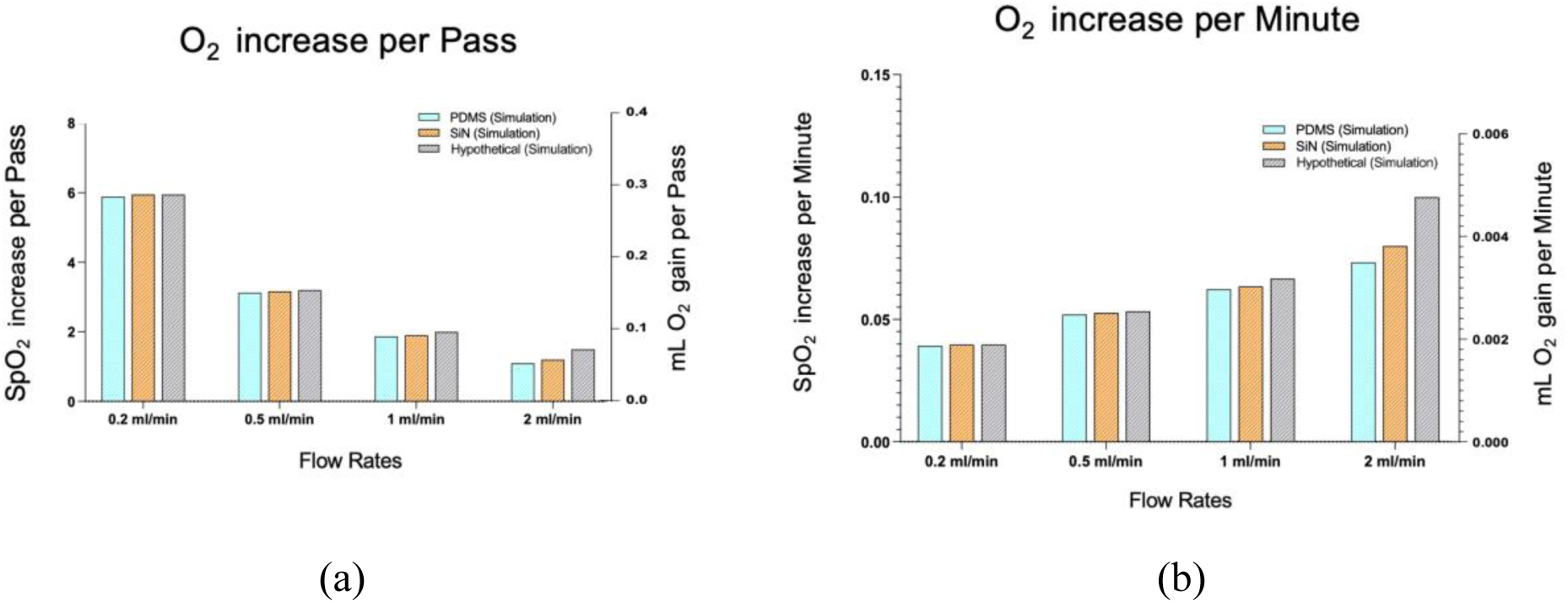
SpO2% rise and ml O_2_ gain of blood for PDMS, nanoporous SiN, and hypothetical membrane at different flow (simulation), (a) per-pass, (b) per minute. In each panel, the left y-axis shows SpO_2_ % rise and the right y-axis shows ml O_2_ gain

## 4. Discussion

In the nanoporous SiN and polypropylene membranes, oxygen transport occurs through a pore-mediated diffusion mechanism. Unlike non-porous PDMS membranes, where gas molecules must diffuse through the solid material, these membranes contain micro-or nano-sized pores that allow oxygen to enter and fill the pore spaces directly. Oxygen molecules then diffuse through these pores without interacting with the membrane material itself. Upon reaching the blood side of the ECMO system, oxygen exits the pores and diffuses directly into the blood. This pore-centric approach enables efficient gas transfer, as the pathway for oxygen diffusion is governed by the pore architecture rather than by the intrinsic diffusivity of the membrane material. In contrast, the PDMS membrane lacks such pores, so oxygen must diffuse directly through the membrane material itself. This happens as oxygen molecules diffuse through the polymer matrix of PDMS by passing through the interconnected network of polymer chains. This occurs because PDMS is an amorphous polymer with a flexible and loosely packed molecular structure. The polymer chains create regions of free volume within the material, allowing small gas molecules, such as oxygen, to diffuse through governed by the concentration gradient across the membrane. The diffusion in PDMS is thus slower, as it depends on the material’s intrinsic permeability to oxygen rather than on a pore-mediated pathway. Fig. 8 shows the intrinsic difference between oxygen molecules passing through nanoporous SiN membrane and PDMS membrane.

**Fig. 8.**
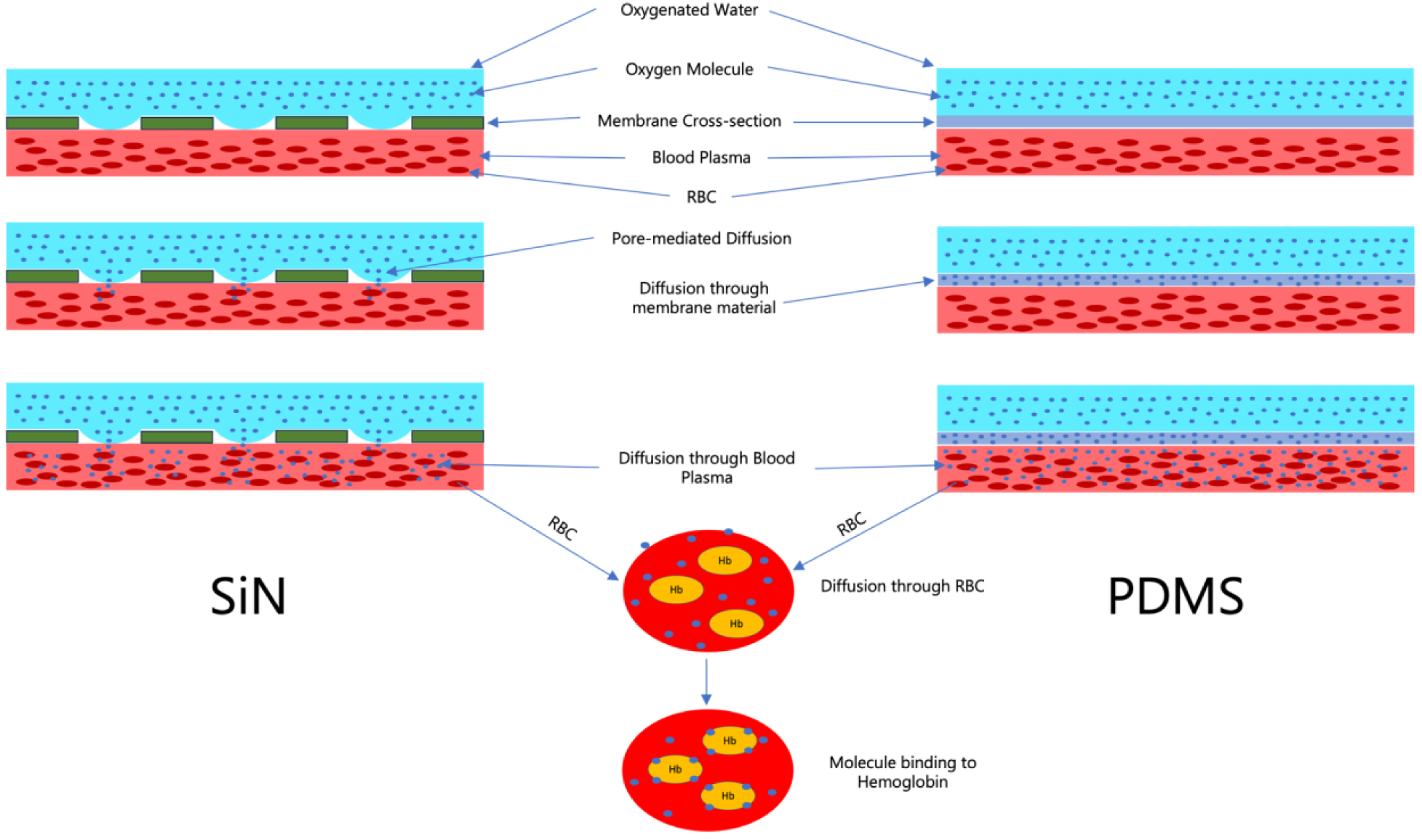
The multilayer diffusion process of an oxygen molecule from the oxygen side to the RBC in the blood side in the microfluidic ECMO for nanoporous SiN and PDMS membrane. Oxygen molecules are transported from the oxygenated water side to the blood side through pore-mediated diffusion for the SiN membrane, and diffusion through membrane material (through the PDMS lattice) for the PDMS membrane

Regardless of membrane properties (Table 1) and mechanistic differences between the membranes, it was observed that the oxygen transport efficiency of different membrane devices was essentially the same. Theoretically, nanoporous SiN, with its ultra-high permeability, was expected to provide the highest oxygen transport efficiency, followed by Polypropylene and PDMS. However, experimental results indicated no significant differences in oxygenation performance across the three membrane types (Fig. 5) (p >0.05). These findings indicate that the membrane material has minimal influence on oxygen transport efficiency in microfluidic ECMO systems. Instead, the rate-limiting steps in oxygen transfer appear to be governed by processes on the blood side. This finding was further supported by the observed oxygen transport performance of a hypothetical infinitely permeable membrane. Even with perfect oxygen transfer efficiency of the membrane, the oxygen increase remained essentially the same as that achieved with the three tested membranes (Fig. 7).

As illustrated in Fig. 5a and 5c, oxygen transport per-pass decreases as flow rate increases due to reduced residence time of red blood cells (RBCs) adjacent to the oxygenating membrane. At lower flow rates, prolonged membrane contact allows more complete oxygen diffusion, with residence times of approximately 47.5, 19, 9.5, and 4.75 seconds at flow rates of 0.2, 0.5, 1, and 2 mL/min, respectively. Higher flow rates reduce equilibration time, diminishing per-pass oxygen gain; however, total oxygen delivery per unit time may increase (Fig. 5b and 5d) as larger blood volumes are processed, compensating for reduced oxygenation per cycle. At excessively high flow rates, gas exchange efficiency declines as RBCs traverse the membrane too quickly, potentially causing total oxygenation to plateau or decrease. For quantification, oxygen gain was reported in “per-pass” terms, providing a flow-independent metric that isolates membrane performance, enabling consistent comparison across flow rates, membrane types, and configurations. This approach allows clinicians and engineers to evaluate and optimize oxygenation efficiency while avoiding under or oversaturation.

The oxygenation process within the ECMO system comprises a series of steps: first, oxygen moves across the membrane (through pore-mediated diffusion or diffusion through non-porous membrane material); then, it dissolves into the blood plasma; subsequently, it diffuses into red blood cells (RBCs) and binds to hemoglobin (Fig. 8). The final binding step to hemoglobin within RBCs is the most critical for achieving elevated blood oxygen levels. This multi-step process is inherently slower than the diffusion across the membrane itself, meaning that oxygen transport within the blood, particularly the binding kinetics to hemoglobin, primarily limits the overall oxygen gain. Factors such as blood kinetics, hemoglobin affinity, RBC membrane permeability, and the concentration of free hemoglobin in the blood collectively determine how much oxygen is transferred into the blood per pass.

The similarity in oxygen transport observed among nanoporous SiN, polypropylene, and PDMS membrane devices suggests that optimizing ECMO oxygenation in microfluidic systems may benefit more from addressing the blood-side limitations rather than from selecting membrane materials solely based on gas permeability. For instance, the introduction of mixing elements in the blood side geometry that enhances mixing, modulation of hemoglobin oxygen affinity, or enhancing hemoglobin concentration could potentially have a greater impact on oxygen transfer efficiency than membrane material changes. Our findings underline the need for a paradigm shift in ECMO design for microfluidic systems, focusing on blood-side enhancement to overcome diffusion and binding limitations.

The results of this study should be considered with an understanding of potential influences from the experimental setup and materials used; however, the findings remain robust within the tested conditions. First, the variability in blood samples, such as donor-to-donor or species-to-species differences in hemoglobin concentration and overall blood composition, could impact oxygen transport performance. To minimize the influence of blood variability, all membranes tested on a given day were evaluated using the same lot of blood. Second, the flow rates selected for the experiments, while representative of microfluidic ECMO applications, may not encompass all physiological or clinical conditions. Higher or lower flow rates could alter RBC residence times and mixing dynamics, potentially affecting oxygen transfer performance. Finally, the geometry of the device, including channel dimensions and membrane placement, may play a critical role in determining flow patterns and oxygen transport. Different geometrical configurations may yield varied oxygenation results, making it essential to consider these design factors when extrapolating the findings to broader ECMO applications. These limitations emphasize the need for additional studies that explore a wider range of flow rates, device geometries, and blood characteristics to generalize the conclusions drawn from this study.

Although membrane material does govern oxygen transfer efficiency in microfluidic ECMO systems in the conditions that we tested, its importance in other applications remains an area of ongoing investigation. For example, in dialysis, efficient removal of waste products like urea relies heavily on the membrane’s ability to facilitate diffusion from the blood side to the dialysate side. Here, urea diffuses across the membrane due to a concentration gradient, moving from the higher concentration in the blood to the lower concentration in the dialysate. Unlike oxygenation in ECMO, dialysis involves minimal biochemical restrictions, as urea does not need to bind to a carrier protein. Instead, the dialysate continuously flushes away urea from the membrane, allowing for effective clearance and maintaining low urea levels in the blood. A seminal study by Hill et al. demonstrated that a nanoporous SiN membrane in a microfluidic dialysis device offers improvements over conventional dialysis membranes for a 300 µl/min flow rate (Hill et al. 2020). In this work it was shown that, nanoporous SiN membranes achieved area-normalized clearance of up to 60,000 mL min^-1^m^-2^ for small solutes—more than 50× higher than the ∼1,000 mL min^-1^m^-2^ typical of commercial polysulfone (PSU) and cellulose triacetate (CTA) membrane dialyzers. In a uremic rat model, 110 mm² of NPN-O lowered serum urea by 26% in 4 hours, whereas conventional membranes with ∼220 mm² area had no measurable effect.

The efficiency of the dialysis process depends not only on membrane permeability but also on the diffusion rate of urea in water (1.382 × 10⁻⁵ cm²/s at 25°C) (Winkelmann 2018), which is critical for urea clearance from the blood. In this process, a constant urea concentration gradient is maintained by the flowing dialysate. This gradient is the sole driving force for urea transport through the dialysis membrane, ensuring its removal from the blood. Although the diffusion rate of O₂ in plasma is similar in magnitude to that of urea in water, blood oxygenation is more complex because multiple biological and chemical factors also influence its overall efficiency.. In the ECMO blood side, oxygen diffuses first through the plasma (with a diffusion rate of 1.62 × 10⁻⁵ cm²/s at 25°C) (Goldstick et al. 1976) and then through the RBC membrane, which has a significantly lower diffusion rate (5 × 10⁻⁸ cm²/s) (Moll 1969). Furthermore, oxygen must bind to hemoglobin within RBCs, making the amount of hemoglobin present and the oxygen-binding affinity of hemoglobin crucial for determining oxygen uptake in blood.

Thus, while membrane material is not the primary factor in blood oxygenation due to the presence of other significant resistances, such as hemoglobin binding and RBC membrane diffusion, it becomes crucial in applications where the membrane itself is the dominant resistance to transport, as seen in dialysis. This distinction highlights the need to consider the relative contributions of various resistances when selecting membrane materials. In cases where membrane resistance is the primary bottleneck, optimizing permeability can lead to substantial improvements. However, in systems like ECMO, where blood-side resistances play a larger role, addressing these factors is likely to have a greater impact on overall efficiency.

## Acknowledgement

The authors would like to thank SimPore Inc. for kindly providing the SiN and PDMS membranes used in this study.

## Author Contributions

N.I.: Conceptualization, Methodology, Formal analysis, Writing - Original Draft, Review & Editing, Visualization. W.A.S.: Validation, CFD analysis, Data Curation, Formal analysis. S.W.D.: Conceptualization, Methodology, Validation, Investigation, Supervision, Resources, Writing - Review & Editing, Supervision, Project administration.

